# Faces and voices in the brain: a modality-general person-identity representation in superior temporal sulcus

**DOI:** 10.1101/338475

**Authors:** Maria Tsantani, Nikolaus Kriegeskorte, Carolyn McGettigan, Lúcia Garrido

## Abstract

Face-selective and voice-selective brain regions have been shown to represent face-identity and voice-identity, respectively. Here we investigated whether there are modality-general person-identity representations in the brain that can be driven by either a face or a voice, and that invariantly represent naturalistically varying face and voice tokens of the same identity. According to two distinct models, such representations could exist either in multimodal brain regions (Campanella and Belin, 2007) or in face-selective brain regions via direct coupling between face- and voice-selective regions (von Kriegstein et al., 2005). To test the predictions of these two models, we used fMRI to measure brain activity patterns elicited by the faces and voices of familiar people in multimodal, face-selective and voice-selective brain regions. We used representational similarity analysis (RSA) to compare the representational geometries of face- and voice-elicited person-identities, and to investigate the degree to which pattern discriminants for pairs of identities generalise from one modality to the other. We found no matching geometries for faces and voices in any brain regions. However, we showed crossmodal generalisation of the pattern discriminants in the multimodal right posterior superior temporal sulcus (rpSTS), suggesting a modality-general person-identity representation in this region. Importantly, the rpSTS showed invariant representations of face- and voice-identities, in that discriminants were trained and tested on independent face videos (different viewpoint, lighting, background) and voice recordings (different vocalizations). Our findings support the Multimodal Processing Model, which proposes that face and voice information is integrated in multimodal brain regions.

**Significance statement:** It is possible to identify a familiar person either by looking at their face or by listening to their voice. Using fMRI and representational similarity analysis (RSA) we show that the right posterior superior sulcus (rpSTS), a multimodal brain region that responds to both faces and voices, contains representations that can distinguish between familiar people independently of whether we are looking at their face or listening to their voice. Crucially, these representations generalised across different particular face videos and voice recordings. Our findings suggest that identity information from visual and auditory processing systems is combined and integrated in the multimodal rpSTS region.

## Introduction

Looking at a familiar person’s face or listening to their voice automatically grants us access to a wealth of information regarding the person’s identity, such as their name, our relationship to them, and memories of previous encounters. Knowledge about how the brain processes faces and voices separately has advanced significantly over the past twenty years: functional magnetic resonance imaging (fMRI) revealed cortical regions that are face-selective (Kanwisher et al., 1997) and regions that are voice-selective (Belin et al., 2000). Recent advances using multivariate classification methods have further shown that some of these regions are important for identification. In particular, face-selective regions in the posterior occipitotemporal lobe, anterior temporal lobe, and posterior superior temporal sulcus (pSTS) can discriminate different facial identities (Kriegeskorte et al., 2007; Nestor et al., 2011; Goesaert and Op de Beeck, 2013; Verosky et al., 2013; Axelrod and Yovel, 2015; Collins et al., 2016; Visconti Di Oleggio Castello et al., 2017). Crucially, a number of studies also found representations in these regions that generalised across different images of the same person (Anzelotti et al., 2014; Anzelotti and Caramazza, 2016; Guntupalli et al., 2017), i.e., were able to “tell people together” (Jenkins et al., 2011; Burton, 2013). Similarly for voices, Formisano et al. (2008) found voice-identity representations in the right STS and Heschl’s gyrus that could both discriminate between speakers and generalise across different vowel sounds spoken by the same voice.

Despite these advances, we still have a limited understanding of how the brain combines and integrates face and voice information. Two major models have been put forward. The *Multimodal Processing Model* proposes that there are multimodal systems that process information about people and receive input from both face- and voice-responsive regions (Ellis et al., 1997; Campanella and Belin, 2007). Patient (Ellis et al., 1989; Gainotti, 2011) and fMRI studies (Shah et al., 2001; Joassin et al., 2011; Watson et al., 2014a) suggest the anterior temporal lobe, the posterior cingulate cortex, the STS, and the hippocampus as candidate multimodal regions. In contrast, the *Coupling of Face and Voice Processing Model* proposes that the direct coupling between face- and voice-responsive brain regions is crucial for the integration of person-identity information (von Kriegstein et al., 2005). In particular, fMRI studies have shown that voice recognition of familiar (or recently learned) people is associated with increased activation in face-responsive regions of the fusiform gyrus (von Kriegstein et al., 2005, 2006, 2008; von Kriegstein and Giraud, 2006).

In this study, we tested the predictions from these two models by investigating whether there are modality-general person-identity representations in multimodal regions (Multimodal Model) and/or in face- and voice-selective regions (Coupling Model). We used representational similarity analysis — RSA (Kriegeskorte et al., 2008a, 2008b) to compare the representational geometries of face- and voice-elicited person-identities, and to investigate the degree to which pattern discriminants for pairs of identities generalise from one modality to the other. We predict that, if a region shows a modality-general person-identity representation, the representational geometry of face and voice identities will match, and/or pattern discriminants will generalise across faces and voices. Two recent studies found some support for the Multimodal Model by showing that multimodal regions in the STS and inferior frontal gyrus (Hasan et al., 2016; Anzelotti and Caramazza, 2017) could discriminate between the activation patterns of two face-identities based on voice information (and vice-versa). However, these studies did not show that the regions that could decode identities across modalities could also decode identities within each modality, which is a crucial feature of modality-general person-identity representations. Furthermore, these studies used very few identities and tokens per identity. In our study, we included multiple, naturalistically varying face videos and voice recordings of 12 different identities. Thus, we were able to sample the variability of visual and auditory appearance that we are exposed to in everyday life, and to better capture processes of person identification, which are distinct from image or sound recognition (Burton, 2013).

## Materials and Methods

### Overview of study

In this study, we measured fMRI activation patterns in response to the faces and voices of 12 famous individuals. It was important to use highly familiar individuals because we needed to guarantee that participants were well acquainted with the faces and voices of those individuals. We thus only recruited participants for the full study if they demonstrated that they were familiar with the majority of the famous individuals in an online Recognition Task. The full study consisted of two MRI scanning sessions and one behavioural session, with each session taking approximately 90 minutes. All three sessions took place on separate days. Before entering the scanner at the start of the first MRI session, participants repeated the Recognition Task in the presence of the experimenter and also completed a Familiarity Task in which they rated all face and voice stimuli on perceived familiarity.

In each MRI session participants completed three functional runs (main experimental runs) in which they viewed the faces and listened to the voices of the famous people in an event-related design. In addition, participants underwent two structural scans (one in each session) and functional localisers for face-selective, voice-selective, and multimodal regions of interest (ROIs). Across both sessions participants completed at least one run (in most cases two) of (1) the temporal voice area (TVA) localiser (Belin et al., 2000), (2) a face localiser, (3) a multimodal (face-voice) localiser, and (4) a voice localiser. Finally, participants completed a behavioural testing session. In this behavioural session they rated the famous faces and voices on various social and perceptual dimensions; however, these results are not included here.

To investigate the existence of modality-general person-identity representations in each of our ROIs we used RSA (Kriegeskorte et al., 2008a, 2008b; Kriegeskorte and Kievit, 2013) to compare the representational geometry of face-identities with the representational geometry of voice-identities (Analysis A), and to investigate the degree to which pattern discriminants for each pair of identities generalise from one modality to the other (Analysis B). Analysis A focused on the representational geometry of all of identities, i.e. the entire structure of pairwise distances between the activity patterns elicited by these identities in each modality, and compared geometries across modalities. Analysis B focused on the discriminability of pairs of identities, and used a linear discriminant computed in one modality to test discriminability of the same pair of identities in the other modality (in a similar way to traditional pattern classification methods). These two analyses complement each other and allowed us to test different predictions regarding the nature of modality-general person-identity representations.

For Analysis A (RSA comparing representational geometries), we predicted that brain regions with modality-general person-identity representations would show matching representational geometries for face-identities and voice-identities. This analysis is constrained by two assumptions. The first assumption is that there is sufficient variability in the representational distances between different identities within-modality, i.e. different degrees of similarity between identities. If all identities are equally distinct from each other, we do not expect to find correlations between geometries across the two modalities. The second assumption is that modality-general information dominates over any modality-specific information that may be present in the same voxels. Specifically, it is possible that the voxels comprising the pattern estimates contain both unimodal and multimodal neurons (Quiroga et al., 2009). In this case, the influence of modality-specific information on the representational distances between all identities could override the influence of modality-general information on the representational geometry, and could result in non-matching representational geometries across modalities.

We thus also conducted Analysis B (RSA investigating identity discriminability), and we predicted that brain regions with modality-general person-identity representations would be able to discriminate between pairs of identities in one modality based on their representational distance in the other modality. This analysis focuses on one pair of identities at a time, and thus is not affected by the degree of variability in the representational distances between all identities. In addition, this analysis is focused on pattern discriminants that generalise across modalities, and therefore we believe that it is more sensitive to detect modality-general person-identity representations even in the presence of modality-specific information.

### Participants

Participants were recruited at Royal Holloway, University of London and Brunel University London to take part in a behavioural and fMRI experiment. All participants were required to be native English speakers aged between 18 and 30, and to have been resident in the UK for a minimum of 10 years. These requirements were set to increase the likelihood of participants being familiar with the famous people whose faces and voices were presented in the experiment. In addition, participants completed an online Recognition Task (see below) as part of the screening procedure for the study and were only invited if they were able to recognise at least 75% of our set of famous people from both their face and their voice.

Thirty-one healthy adult participants were recruited who matched all the above criteria. One participant was excluded from the study after the first MRI session due to excessive head movement in the scanner (more than 3 mm in any direction within one run). The final sample consisted of 30 participants (eight males) with mean age of 21.2 years (SD=2.37, range=19-27). All reported normal or corrected-to-normal vision and normal hearing, provided written informed consent and were reimbursed for their participation. The study was approved by the Ethics Committee of Brunel University London.

### Recognition Task

Participants completed a face and voice Recognition Task to determine whether they could recognise at least 75% of the famous people (i.e. at least 9 out of 12) from both the face and the voice. Face stimuli consisted of single photographs of each of the 12 famous people that were obtained from the Internet through Google Image searches. Photographs included the top part of the body and were front-facing. Voice stimuli consisted of single sound-clips for each of the 12 famous people and were obtained from YouTube videos. Sound-clips were approximately 8-seconds long and were root-mean-square (RMS) normalized using Praat (version 5.3.80; Boersma and Weenink, 2014; www.praat.org). None of these face or voice stimuli were presented in the main experiment.

Stimuli were presented using Qualtrics (Qualtrics, Provo, UT). For each stimulus participants had to identify the person shown in the picture or the person speaking (by providing their name or other uniquely identifying biographical information). In the online task participants typed their responses below each stimulus, and in the lab task responses were made verbally.

### Stimuli for Familiarity Task and Main Experimental Runs

Six silent, non-speaking video clips of moving faces, and six sound clips of voices for each of the 12 famous people (six female, six male) were obtained from videos on YouTube (in total, 72 stimuli per modality). These people had been identified in our pilot studies as having highly recognisable faces and voices within samples of native English speakers between the ages of 18-30 who have been resident in the UK for a minimum of 10 years. This list of famous people included actors, pop stars, politicians, comedians, and TV personalities: Alan Carr, Beyonce Knowles, Daniel Radcliffe, Emma Watson, Arnold Schwarzenegger, Barack Obama, Sharon Osbourne, Kylie Minogue, Graham Norton, Cheryl Cole, Barbara Windsor, and Jonathan Ross.

The face stimuli were selected so that the background did not provide any cues to the identity of the person. Other than the absence of speech, there were no constraints on the type of face movement. Examples of face movements included nodding, smiling, and rotating the head. However, all stimuli were selected to be primarily front-facing. Face stimuli were edited using Final Cut Pro X (Apple, Inc.) so that they were three seconds long and centred on the bridge of the nose. Six video-clips of the face of the same person were obtained from different original videos set in a different background.

Voice stimuli were edited using Audacity® 2.0.5 recording and editing software (RRID:SCR_007198) so that they contained three seconds of speech after removing long periods of silence. Voice stimuli were converted to mono with a sampling rate of 44100, low-pass filtered at 10KHz, and RMS normalised using Praat. Six sound clips of the voice of the same person were obtained from different original videos. All of the voice stimuli had a different verbal content and were non-overlapping. The stimuli were selected so that the speakers’ identity could not be determined based on the verbal content, conforming to the standards set by Van Lancker et al. (1985) and Schweinberger et al. (1997).

### Familiarity Task

Before entering the scanner, participants rated all stimuli that would be presented in the main experimental runs on perceived familiarity. Participants were presented with the face stimuli first, followed by the voice stimuli, in separate blocks. Stimuli were presented using the Psychophysics Toolbox (version 3; RRID:SCR_002881; Brainard, 1997; Pelli, 1997) running in Matlab (version R2013b; MathWorks; RRID:SCR_001622). Face stimuli were presented in the centre of the screen. Participants listened to the voice stimuli through headphones (Sennheiser HD 202). Participants rated each stimulus on scale from 1 (very unfamiliar) to 7 (very familiar). Each block took approximately 5 minutes to complete.

### MRI data acquisition

Participants were scanned using a 3.0 Tesla Tim Trio MRI scanner (Siemens, Erlangen) with a 32-channel head coil at the Combined Universities Brain Imaging Centre (CUBIC) at Royal Holloway, University of London. In each of the two scanning sessions, a whole-brain T1-weighted anatomical scan was acquired using magnetization-prepared rapid acquisition gradient echo (MPRAGE) [1.0 x 1.0 in-plane resolution; slice thickness, 1.0mm; 176 axial interleaved slices; PAT, Factor 2; PAT mode, GRAPPA (GeneRalized Autocalibrating Partially Parallel Acquisitions); repetition time (TR), 1900ms; echo time (TE), 3.03ms; flip angle, 11°; matrix, 256x256; field of view (FOV), 256mm].

For all functional runs T2*-weighted whole-brain functional scans were acquired using echo-planar imaging (EPI) [3.0 x 3.0 in-plane resolution; slice thickness, 3.0mm; PAT, Factor 2; PAT mode, GRAPPA (GeneRalized Autocalibrating Partially Parallel Acquisitions); 34 sequential (descending) slices; repetition time (TR), 2000ms; echo time (TE), 30ms; flip angle, 78°; matrix, 64x64; field of view (FOV), 192mm]. For the majority of participants, slices covered all parts of the brain except for the most dorsal part of parietal cortex. In each experimental run we obtained 293 brain volumes, in the TVA localiser we obtained 251 brain volumes, and in each run of the face, voice, and multimodal localiser runs we obtained 227 brain volumes.

### fMRI data pre-processing

Data were pre-processed using Statistical Parametric Mapping (SPM12; Wellcome Department of Imaging Science, London, UK; RRID:SCR_007037; http://www.fil.ion.ucl.ac.uk/spm) operating in Matlab. Pre-processing was performed separately for each scanning session. All runs within each session (main experiment or localizer runs) were pre-processed together. The first three EPI images in each run (dummy scans) were discarded to allow for T1-equilibration effects. Images were slice-time corrected based on the middle slice in each volume and then realigned to correct for head movement based on the first image. The structural image in native space was then coregistered with the realigned mean functional image and segmented into grey matter, white matter, and cerebrospinal fluid. No smoothing was performed on the images from the experimental runs. Functional images from the localiser runs were smoothed with a 4-mm Gaussian kernel (full width at half maximum).

After separate pre-processing of the images in each session, images from the second scanning session were realigned to the structural image from the first session. Specifically, the structural image from session two was coregistered to the structural image from session one, and the transformation was then applied to all functional images from session two. As a result, all functional images were in the same space.

### Functional localiser runs

#### TVA localiser

We used the TVA localiser developed by Belin et al. (2000) which contains vocal and non-vocal auditory stimuli. Stimuli were presented in 40 blocks of 8 seconds each. Vocal stimuli were presented in 20 blocks and included speech and non-speech vocalisations obtained from 47 speakers (Pernet et al., 2015). Non-vocal stimuli were presented in 20 blocks and consisted of industrial sounds, environmental sounds, and animal vocalisations. Within each block stimuli were presented in a random order that was fixed across participants. Participants were instructed to close their eyes and focus on the sounds. The TVA localiser was presented directly after the main experimental runs. The duration of a single run was approximately 10 minutes.

#### Face, Voice, and Multimodal localisers

We created new face, multimodal, and voice localiser runs that shared the same experimental design and presented stimuli from comparable categories (people and objects/scenes). Importantly, we used videos and not static images of faces. Dynamic face stimuli have been shown to be more effective that static face stimuli for localising face-selective regions (Fox et al., 2009; Pitcher et al., 2011). Stimuli used for the face localiser were silent, non-speaking video clips of famous and non-famous (French celebrities unknown to our participants) moving faces, and silent video clips of moving large objects and natural or manmade visual scenes (such as videos of airplanes, trains, traffic, rainforests, waves on a beach) obtained from videos on YouTube. For the multimodal localiser the stimuli were audio-visual and included videos clips of the faces of famous and non-famous people speaking, and video clips of moving large objects and natural or manmade scenes (same categories as above). For the voice localiser we presented voice clips of famous and non-famous people, and sound clips of manmade or natural environmental sounds (same categories as used in the other two types of localisers), with no video.

Videos (640 x 360 pixels) were presented at the centre of the screen. The screen resolution was 1024 x 768 pixels, and from a distance of 85 cm, the videos subtended 20.83 x 12.27 degrees of visual angle. Audio stimuli were presented via MR-compatible earbuds (S14; Sensimetrics Corp.), which participants used for each entire scanning session. Each stimulus lasted 8 seconds and each run presented 48 stimuli. Stimuli were presented in pairs (24 pairs) showing the same person (such as two videos of Brad Pitt) or the same category of objects or scenes (such as two videos of trains). Eight pairs showed stimuli from famous people, eight pairs showed stimuli from non-famous people, and eight pairs showed object/scene stimuli. Participants were encouraged to always fixate at the centre of the screen. Participants performed a one-back task in which they had to detect the exact same stimulus repetition within each pair, which occurred in approximately 15% of the trials. A 16-second period of fixation was presented at the end of each run and twice in the middle of each run (every 16 trials).

The order of the face, voice, and multimodal localisers was counterbalanced across participants. For participants who completed two runs of each localiser, different identities were presented on each run. The duration of each localiser run was approximately 8 minutes.

#### General linear models

To identify face-selective (face localiser), voice-selective (voice localiser and TVA localiser), and people-selective (multimodal localiser) brain regions, we computed mass univariate time-series models for each participant. Regressors modelled the blood-oxygenation-level-dependent (BOLD) response following the onset of the stimuli and were convolved with a canonical hemodynamic response function (HRF). We also used a high-pass filter cutoff of 128 seconds, and autoregressive AR(1) model to account for serial correlations. For the face, voice, and multimodal localisers there were three experimental regressors: (1) famous faces/voices/people, (2) non-famous faces/voices/people, and (3) objects and scenes. For the TVA localiser there were two experimental regressors: (1) voices and (2) non-voices. For all localisers six head motion parameters computed during realignment were included as covariates. Selectivity was defined with a *t*-test contrasting the responses to faces/voices/people (famous and non-famous) *versus* responses to the control stimuli.

#### ROI definition

We used probabilistic maps from previous studies to define regional masks in which we predicted that our regions of interest (ROIs) would be located. We then defined ROIs by extracting all selective voxels within those regional masks for each participant. This approach is similar to the one implemented by Julian et al. (2012) and avoids experimenter biases in ROI definition.

Probabilistic maps were thresholded to only show voxels that were present in 20% of the participants and binarised to create regional masks. We used a probabilistic map of the TVAs created by Pernet et al. (2015) and obtained from neurovault (http://neurovault.org/images/106/) to create separate masks for the right and left TVA (rTVA, lTVA). For all other regional masks, we used probabilistic maps that were obtained from a previous study conducted in the lab (unpublished data). In this previous study we tested 22 participants using the same face and voice localisers as the current study (we did not use the multimodal localiser in this previous study). We defined face-selective and voice-selective *t*-test images for each participant, thresholded each image at *p*<.05 (uncorrected), binarised the resulting image, and summed all images across participants to create face-selective and voice-selective probabilistic maps. In cases where there was some overlap between the masks for different regions we manually defined the borders of these masks using anatomical landmarks.

Regional masks of face-selective regions were created for the right fusiform face area (rFFA), the right occipital face area (rOFA), and the right posterior superior temporal sulcus (rpSTS). Regional masks of voice-selective regions were created for the right and the left superior temporal sulcus and gyrus (rSTS/STG, lSTS/STG). Regional masks of multimodal regions were created based on joint face-selective and voice-selective probabilistic maps. These masks were created for a number of regions that showed *both* face-selective and voice-selective responses in most participants: precuneus/posterior cingulate, orbitofrontal cortex (OFC), frontal pole (FP), and right and left temporal pole with anterior inferior temporal cortex (rTP-aIT, lTP-aIT) — we considered the TP and aIT together as the peaks were difficult to separate in most participants. We did not create a mask of the multimodal STS using this method due to the voice-selective STS region being much larger than the face-selective STS region. However, there was large overlap between our mask of the face-selective rpSTS and our masks of the rSTS/STG and rTVA, suggesting that this face-selective rpSTS region also responds to voices.

All of the regional masks (in MNI space) were registered and resliced to each participant’s native space using FSL (version 5.0.9; RRID:SCR_002823; Jenkinson et al., 2012). These masks were then used to extract ROIs from the *t*-test maps obtained from the contrasts of interest from the face, voice, TVA, and multimodal localisers from the current study. All voxels that fell within the boundaries of the mask and that were significantly activated at *p*<.001 (uncorrected) were included in the subject-specific ROI. If there were fewer than 30 voxels at *p*<.001 the threshold was lowered to *p*<.01 or *p*<.05. If we could not define 30 selective voxels even at *p*<.05, the ROI for that participant was not included in the analyses. We required that all ROIs be present in at least 20 participants (out of 30).

### Main experimental runs: Experimental design and statistical analysis

#### Design and procedure

Face and voice stimuli were presented using the Psychophysics Toolbox via a computer interface inside the scanner. Face and voice clips of all 12 identities were intermixed within each run. A fixation point was always present and participants were asked to fixate. The videos were 640 x 360 pixels and, from a viewing distance of 85cm, videos subtended 20.83 x 12.27 degrees of visual angle. The six face videos and the six voice recordings for each of the 12 identities were evenly distributed among the three runs so that each run contained two different videos of the face and two different recordings of the voice of each identity. Each individual stimulus was presented twice within each run. Therefore, in each run there were 96 experimental trails (48 face trials, 48 voice trials) in total.

Participants performed an anomaly detection task that involved pressing a button when they saw or heard a novel famous person that was not part of the set of the 12 famous people that they had been familiarised with prior to entering the scanner. Therefore, each run also contained 12 task trials presenting six famous faces and six famous voices that were not part of the set of famous people that the participants had been familiarised with.

Stimuli were presented in a pseudorandom order that ensured that within each modality each identity could not be preceded or succeeded by one of the other identities more than once, and that each stimulus could not be succeeded by a repetition of the exact same stimulus. Face and voice clips were presented for three seconds with a SOA of four seconds. Thirty-six null fixation trials were added to each run (~25% of the total number of trials). Thus, each run contained 144 trials in total and lasted approximately 10 minutes.

The presentation order of the three runs was counterbalanced across participants. The same three runs with the same face videos and voice recordings that were presented in scanning session one were also presented in session two. However, the three runs were presented in different orders in both sessions (counterbalanced across participants) and stimuli within each run were presented in a new pseudorandom sequence. As an exception, the stimuli for the task trials were different in the two sessions in order to maintain their novelty.

#### General linear models

We computed mass univariate time-series models for each participant. Models were defined separately for each scanning session and each experimental run (six runs in total). Regressors modelled the BOLD response following the onset of the stimuli and were convolved with a canonical hemodynamic response function (HRF). We also used a high-pass filter cutoff of 128 seconds and autoregressive AR(1) model to account for serial correlations. The 12 different identities in each modality were entered as separate regressors in the model (i.e. 24 regressors). Each of these regressors included the two different face videos and voice recordings of each identity that were presented in the run, as well as the two repetitions of each stimulus. Task trials and six head motion parameters computed during realignment were included as regressors of no interest.

As part of the crossvalidation procedure used in the RSA analyses described below, separate models were estimated for each partition of each crossvalidation fold, thus resulting in parameter estimates and residual time courses for every possible independent partition. For partitions with two runs, data was concatenated before estimating the model. In the analyses described below we used the beta estimates computed at each voxel of each ROI for each of the 24 experimental conditions (12 face-identities and 12 voice-identities).

#### Mean response to faces and voices in ROIs

We conducted an analysis to characterise the responses to faces and voices in each ROI, and to confirm that each ROI showed the expected responsivity to faces and voices. For this analysis, we calculated the mean (across all voxels in each ROI, and across all runs) of the parameter estimates for the 12 face-identities and the mean of the parameter estimates for the 12 voice-identities. For each ROI we tested whether the mean for faces and mean for voices were significantly different from zero (across participants) using one-sample *t*-tests. P values were corrected for 24 comparisons (2 tests x 12 ROIs) controlling the false discovery rate (FDR), with *q*<.05. We also compared the mean for faces with the mean for voices in each ROI using paired *t*-tests. P values were corrected for multiple comparisons (12 comparisons) using FDR with *q*<.05.

#### Analysis A: RSA comparing representational geometries

For this analysis we computed representational dissimilarity matrices (RDMs) for faces and voices (12x12 matrices) separately for each participant, each scanning session and each ROI. We then computed the correlations between pairs of these RDMs (for an example, see Figure 4). These analyses were performed using in-house Matlab code and the RSA toolbox (Nili et al., 2014). To compute the RDMs we used the linear discriminant contrast (LDC), a crossvalidated distance measure (Nili et al., 2014; Walther et al., 2016). For each ROI, each modality (i.e. faces and voices separately), and each scanning session, we calculated the LDC between the pattern estimates (beta estimates across all voxels within an ROI) elicited by the different identities. The resulting 12x12 matrices were symmetric around a diagonal of zeros (Figure 4). Each cell in the RDMs showed the discriminability of the pattern estimates corresponding to a pair of identities in the chosen modality and ROI.

RDMs were computed using leave-one-run-out crossvalidation across the three runs in each session (each run presented the same identities with different stimuli). In each of three crossvalidation folds, the pattern estimates for each identity were computed with data from two runs (partition one) and separately from the pattern estimates from the remaining run (partition two). The pattern estimates from each pair of identities from partition one were used to obtain a linear discriminant, which was then applied to differentiate the activity patterns of the same identity pairs in partition two (Nili et al., 2014; Walther et al., 2016). We applied multivariate noise normalisation by computing a noise variance-covariance matrix based on the residual time courses obtained from the model that was estimated with data from partition one. More specifically, to compute the LDC for each pair of identities we first multiplied the contrast between the patterns of a pair of identities in partition one (the discriminant weights) by the inverse of the noise variance-covariance matrix (after regularisation using the optimal shrinkage method: Ledoit and Wolf, 2004), and transformed the resulting weights to unit length. We then computed the dot product between the resulting vector and the vector with the contrast between the patterns of the same pair of identities from partition two (Carlin and Kriegeskorte, 2017), which resulted in a single value showing the discriminability of those identities. Finally, the resulting RDMs with LDC values from each crossvalidation fold were averaged to create one RDM per scanning session. This procedure resulted in four RDMs per participant per ROI: faces session 1, voices session 1, faces session 2, and voices session 2 (Figure 4). Crossvalidating across runs with different videos of the face and recordings of the voice of each identity ensured that the resulting RDMs represented face- and *voice-identity,* rather than specific face videos and voice recordings.

In order to compare the representational geometries of the face- and voice-identities, the RDMs for each participant were compared across the two scanning sessions using Pearson’s correlation coefficient (Figure 4). We also compared the representational geometries of face and voice-identities within modality across two scanning sessions in order to investigate the stability of the representational geometries across the two scanning sessions. For the *crossmodal comparisons* we compared the face and voice RDMs from session one with the RDMs of the *other* modality in session two (i.e. faces session 1 vs. voices session 2, and voices session 1 vs. faces session 2). For the *unimodal comparisons* we compared the face and voice RDMs from session one with RDMs of the *same* modality in session two (i.e. faces session 1 vs. faces session 2 and voices session 1 vs. voices session 2). At the group level for each ROI we compared the single-subject correlations for each of the four comparisons (two crossmodal, two unimodal) against zero using one-sample one-tailed Wilcoxon signed-rank tests (because correlations are not normally distributed). P values were corrected for multiple comparisons (48 comparisons: 4 tests × 12 ROIs) controlling for FDR with *q* < .05.

#### Analysis B: RSA investigating identity discriminability

For this analysis we computed crossmodal RDMs separately for each participant, each scanning session and each ROI. We used the activity patterns of identity pairs in one modality to create a linear discriminant and then applied the discriminant to the activity patterns of the same identity pairs in the other modality. With this exception, the crossvalidation procedure was identical to the procedure for creating face and voice RDMs for the previous analysis. Two crossmodal RDMs for each ROI were computed using this method: one by applying a linear discriminant based on face data to voice data, and one by applying a linear discriminant based on voice data to face data. The LDC provides a continuous measure of discriminability for each pair of stimuli (Nili et al., 2014; Walther et al., 2016; Carlin and Kriegeskorte, 2017). Importantly, under the null hypothesis the LDC is symmetrically distributed around zero, and thus unbiased. By calculating the mean LDC value across all cells in an RDM for a certain ROI we can determine the overall ability of that ROI to discriminate between identities. Mean LDC values for all participants can then be subjected to random-effects inference comparing against zero. Therefore, we predicted that crossmodal RDMs for regions with modality-general person-identity representations would show mean LDC values that are significantly greater than zero.

In addition to investigating identity discrimination *across modalities* using crossmodal RDMs, we also investigated the ability of each ROI to discriminate between identities *within modality,* using the face and voice RDMs that were created in the previous analysis. We predicted that face or voice RDMs for regions that represent face or voice identity, respectively, would show mean LDC values that are significantly greater than zero.

For this analysis the corresponding RDMs (e.g. faces session 1 and faces session 2) for each scanning session were averaged across the two sessions, and then the mean LDC across the vectorised matrix was calculated. Thus, for each participant and each ROI we obtained four mean LDC values representing (1) face discriminability, (2) voice discriminability, (3a) crossmodal discriminability - face discriminant generalised to voices, and (3b) crossmodal discriminability - voice discriminant generalised to faces. For each ROI and each type of discriminability we entered participants’ LDC values into a one-sample one-tailed *t*-test comparing them against zero. P values were corrected for all comparisons (48 comparisons: 4 tests x 12 ROIs) controlling for FDR with *q*<.05.

#### Code and data accessibility

All the code and data for the above analyses will be made available after publication.

#### Exploratory whole-brain searchlight analyses

Despite including a broad range of functionally defined ROIs, it is possible that modality-general person-identity representations may exist in brain regions not included in our ROIs. Specifically, these representations may exist in brain regions that are not face-selective or voice-selective. Therefore, we used an exploratory whole-brain searchlight analysis to identify potential brain regions with person-identity representations using the same methods as in our main ROI analyses. We note that we focused solely on modality-general person-identity representations in this exploratory analysis, as that was the main aim of this study.

For each participant we created 6mm radius spheres centred on each voxel within a grey-matter mask of their brain (obtained from the segmentation procedure) using the RSA toolbox (Nili et al., 2014) in Matlab. A 6 mm radius resulted in a searchlight sphere of 33 voxels, which matched our requirement for minimum ROI size of 30 voxels in the main analyses. For the analysis comparing representational geometries we computed a face and a voice RDM in each searchlight sphere, averaging the RDMs from both scanning sessions, and then calculated the Pearson correlation between them. Correlations were Fisher z-transformed. The output of this analysis was a whole-brain map of Fisher-transformed correlation coefficients for each participant. For the second analysis investigating identity discriminability we computed a single crossmodal RDM in each searchlight sphere by averaging the crossmodal face-voice RDM with the crossmodal voice-face RDM, and then calculating the mean LDC across the resulting matrix in vector form. The output for each participant was a whole-brain map of mean LDC values.

The whole-brain searchlight maps from each analysis were normalised to MNI space using the normalisation parameters generated during the segmentation procedure and spatially smoothed with 9-mm Gaussian kernel (full width at half maximum) to correct for errors in intersubject alignment. For group-level analysis, all searchlight maps were entered into a one-sample *t*-test to determine whether the correlation coefficient/mean LDC value was significantly greater than zero at each voxel. We used the randomise tool (Winkler et al., 2014) in FSL for inference on the resulting statistical maps (5,000 sign-flips). Clusters were identified with threshold-free cluster enhancement (TFCE), and p-values were corrected for multiple comparisons (FWE < 0.05).

## Results

### Familiarity ratings

Familiarity ratings of both faces and voices were high (Faces: *M* = 6.28, *SD* = 0.5; Voices: *M* = 6.2, *SD* = 0.49). Average familiarity of each identity’s face and voice are shown in Table 1.

**Table 1.** Familiarity ratings of the face and voice of each identity averaged across participants showing the mean (*M*) rating of the face and voice of each identity across all face videos and all voice recordings of that identity, the standard deviation (*SD*) of participants’ ratings of each identity, and the range of mean ratings for the six face tokens and six voice tokens for each identity. The rating scale ranged from 1 (very unfamiliar) to 7 (very familiar).

|  |  | AC | AS | BO | DR | GN | JR | BK | BW | CC | EW | KM | SO |
| --- | --- | --- | --- | --- | --- | --- | --- | --- | --- | --- | --- | --- | --- |
| Faces | M | 6.48 | 5.72 | 6.94 | 6.76 | 6.27 | 6.42 | 6.36 | 5.65 | 6.49 | 6.73 | 5.25 | 6.32 |
|  | SD | 0.75 | 1.19 | 0.2 | 0.59 | 0.85 | 0.59 | 0.94 | 1.45 | 0.79 | 0.45 | 1.48 | 0.92 |
|  | Token | 6.37- | 4.83- | 6.90- | 6.67- | 5.87- | 6.17- | 6.07- | 5.33- | 6.37- | 6.47- | 4.6- | 6.17- |
|  | range | 6.60 | 6.13 | 7 | 6.87 | 6.5 | 6.57 | 6.57 | 5.9 | 6.60 | 6.9 | 5.57 | 6.47 |
| Voices | M | 6.59 | 5.66 | 6.73 | 6.69 | 6.37 | 6.54 | 6.23 | 5.54 | 6.63 | 6.07 | 5.3 | 6.02 |
|  | SD | 0.54 | 1.48 | 0.63 | 0.57 | 0.77 | 0.71 | 1.04 | 1.74 | 0.74 | 0.94 | 1.45 | 1.03 |
|  | Token | 6.37- | 5.43- | 6.7- | 6.57- | 6.07- | 6.3 | 6.07- | 5.47- | 6.4- | 4.6- | 5- | 5.5- |
|  | range | 6.83 | 5.87 | 6.8 | 6.8 | 6.67 | 6.7 | 6.37 | 5.67 | 6.77 | 6.53 | 5.53 | 6.43 |

### ROI definition

Using functional localisers we defined face-selective ROIs (rFFA, rOFA, rpSTS), voice-selective ROIs (rSTS/STG, rTVA, lSTS/STG, lTVA), and multimodal ROIs (OFC, FP, rTP-alT, lTP-aIT, Prec./P.Cing. [including the retrosplenial cortex]) in each participant. We were able to localise these ROIs with at least 30 voxels in all 30 participants, except for the face-selective rFFA (28 participants) and rOFA (29 participants), the Prec./P.Cing. (26 participants), and the OFC (21 participants). We note that the voice-selective ROIs in the right hemisphere (rTVA, rSTS/STG) overlap with each other and with the face-selective rpSTS and the multimodal rTP-aIT ROIs. In addition, the voice-selective ROIs in the left hemisphere (lTVA, lSTS/STG) overlap with each other and with the multimodal lTP_aIT ROI. For visualisation purposes only, probabilistic maps of all ROIs were created by normalising the single subject ROIs to MNI space and summing them. Figure 1 shows those maps thresholded to display all voxels that were present in at least 20% of the participants.

**Figure 1:**
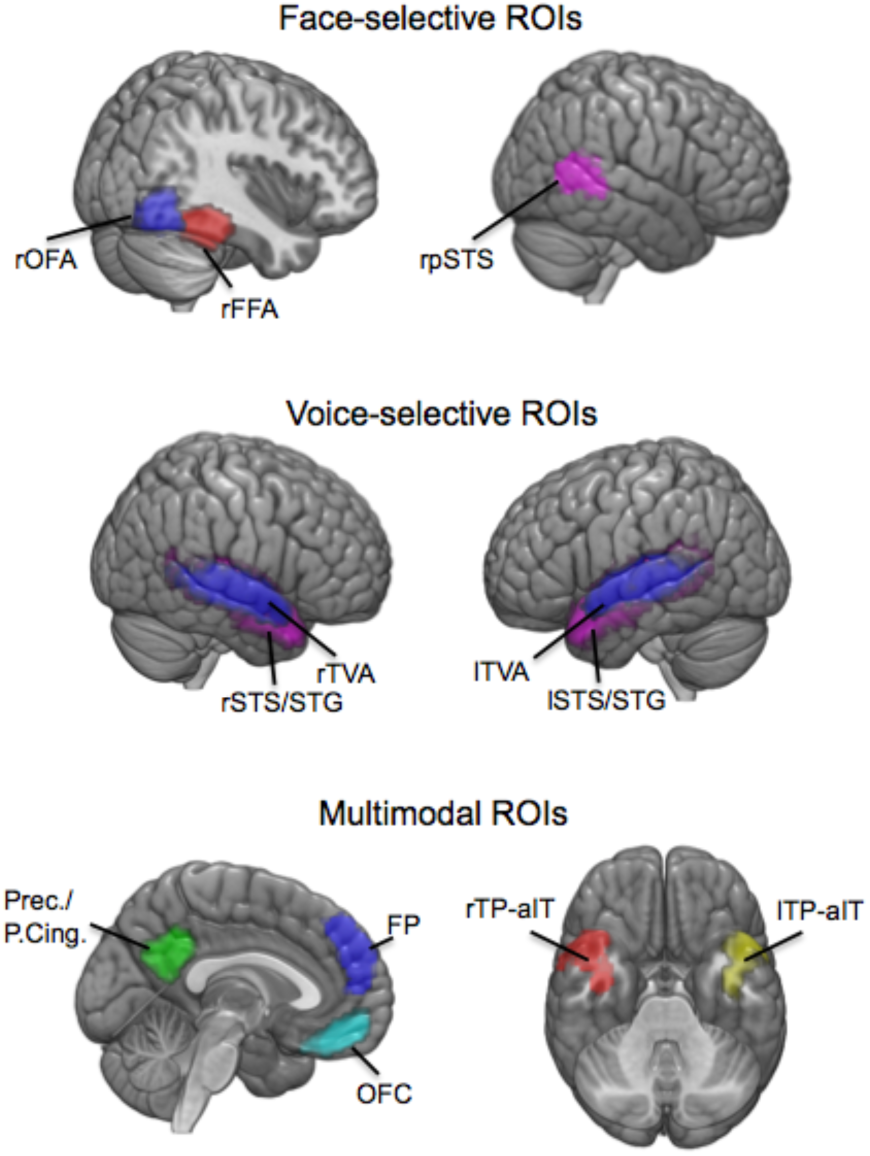
Face-selective, voice-selective, and multimodal ROIs. Location of ROIs that resulted from the face, voice, and multimodal localisers in MNI space. r = right, l = left, FFA = fusiform face area, OFA = occipital face area, pSTS = posterior superior temporal sulcus, STS/STG = superior temporal sulcus/superior temporal gyrus, TVA = temporal voice area, OFC = orbitofrontal cortex, FP = frontal pole, TP = temporal pole, aIT = anterior inferior temporal cortex, Prec = precuneus, P.Cing. = posterior cingulate.

### Mean response to faces and voices in ROIs

In order to confirm that each ROI showed the expected responsiveness to faces and voices, we computed the regional mean of the parameter estimates for faces and for voices across participants for each ROI and modality (Figure 2). As expected, mean beta values for faces were high and significantly greater than zero in all three face-selective ROIs (all one-sample *t*-tests with *p*<.0001). Mean beta values for voices were also significantly greater than zero in the rFFA (*p*<.0001) and rpSTS (*p*<.0001), but not in the rOFA. The rFFA and the rOFA showed significantly greater responses to faces compared with voices (both paired-samples *t*-tests with *p*<.0001). In contrast, the rpSTS showed significantly greater responses to voices compared with faces (*p*=.0002) despite being defined using our face localiser. This is most likely due to the large overlap between this ROI and the voice-selective rSTS/STG and rTVA ROIs. This finding demonstrates that the rpSTS also showed substantial responses to voices.

**Figure 2:**
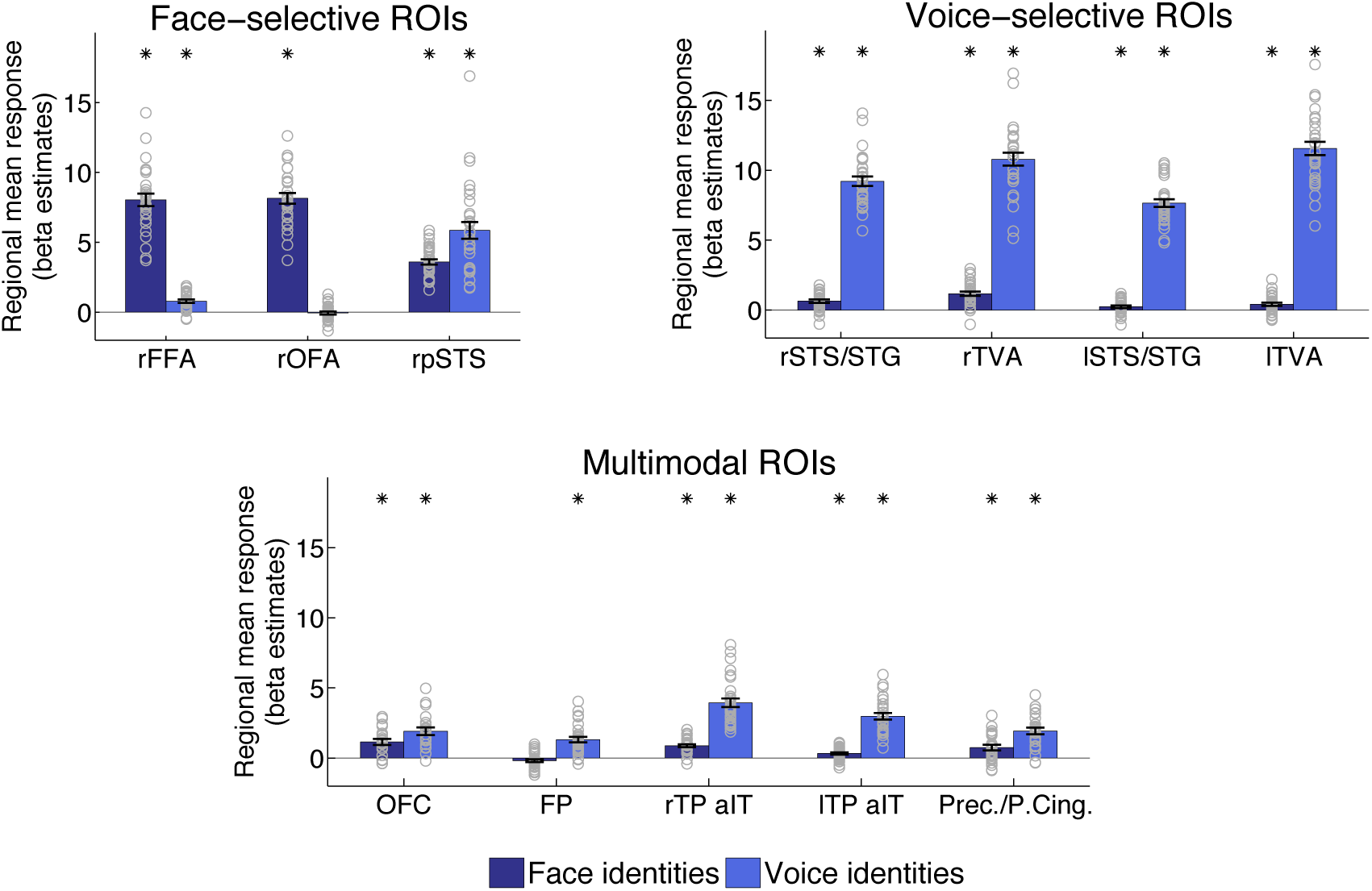
Regional mean responses to faces and voices in ROIs. Regional mean responses for all face-identities and for all voice-identities in face-selective, voice-selective, and multimodal ROIs (mean beta estimates across all voxels of each ROI, and across all runs). Bars show mean responses across participants, error bars show standard error, and grey circles show individual participants. We tested whether mean responses were significantly greater than zero using one-sample *t*-tests across all 30 participants, and stars show significant results at *p*≤.0209 (FDR corrected for all 24 comparisons). We also tested whether mean beta values for faces were significantly different from mean beta values for voices in each ROI using paired *t*-tests across all participants. In all ROIs mean beta values for faces and voices were significantly different at *p*≤.0011 (FDR corrected for all 12 ROIs).

It could be that the responses to voices in rpSTS were due to the voices being familiar, and not because of being voices *per se.* To determine whether this region responded to voices more generally or just to familiar voices, we investigated the responses in rpSTS to familiar voices, unfamiliar voices, and non-voices during the functional voice localisers. For each participant, we calculated the mean parameter estimates across all voxels of the face-selective rpSTS for each condition of the voice localiser (familiar voices, unfamiliar voices, and auditory scenes) and of the TVA localiser (vocal and non-vocal sounds). For the voice localiser, both the familiar and the unfamiliar voices had significantly higher parameter estimates than the auditory scenes (both *p*<.0001). For the TVA localiser, the rpSTS also showed significantly higher responses to voices than non-voices (*p*<.0001). These results show that the face-selective rpSTS also responds to voices in general and not only familiar voices (for similar results, see Deen et al., 2015), and therefore in the rest of this article we will refer to this rpSTS region as displaying multimodal responses.

Returning to the analysis of the parameter estimates for faces and voices during the main experimental runs, the mean beta values for voices were significantly greater than zero for all four voice selective ROIs (all *p<*.0001). Mean beta values for faces were also significantly greater than zero for all voice-selective ROIs (all *p*≤.0209), but the parameter estimates were significantly lower than for voices (all *p*<.0001).

For the multimodal ROIs mean beta values for faces and for voices were significantly greater than zero in all ROIs (all *p*≤.0009) except the frontal pole for faces. This result demonstrates that, although we still included the frontal pole ROI in the main analyses, we cannot be confident about the multimodal responses of this ROI. Also, we note that in all multimodal ROIs (OFC, FP, rTP-aIT, lTP-aIT, Prec./P.Cing.) mean beta values for voices were significantly higher than mean beta values for faces (all *p*≤.0011). We observed this consistently across all participants.

### Analysis A: RSA comparing representational geometries

Our first main analysis compared the representational geometry of the 12 famous identities across and within modalities in each ROI. We computed face and voice RDMs separately for each session using the LDC and compared the RDMs using Pearson correlation (Figures 3 & 4). We then tested whether these correlations were significantly above zero.

**Figure 3:**
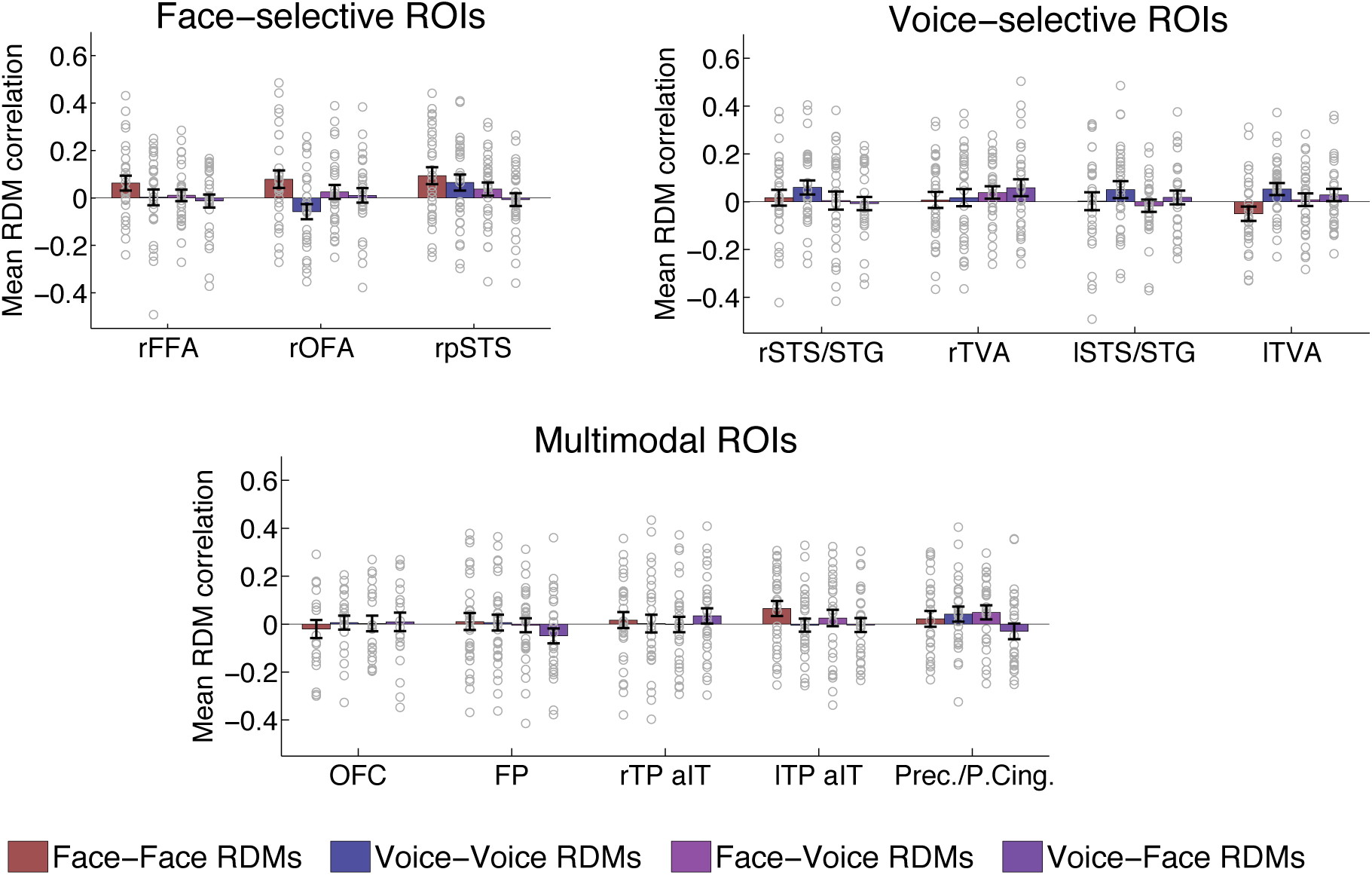
Results of RSA comparing representational geometries. Comparisons between the representational distance matrices (RDMs) from two scanning sessions using Pearson’s correlation coefficient. Bars show mean correlations across participants, error bars show standard error, and grey circles show the correlations of individual participants. Correlations were calculated across scanning sessions and compared face RDMs, voice RDMs, face and voice RDMs, and voice and face RDMs in face-selective, voice-selective, and multimodal ROIs. We tested whether correlations were significantly greater than zero using Wilcoxon signed-rank tests across all 30 participants. No correlations were significant after correction for multiple comparisons at *p*≤.0001 (FDR corrected for all 48 comparisons). Note that in this figure the rpSTS is classed as a face-selective ROI for consistency purposes only, but in fact it demonstrated multimodal properties.

**Figure 4:**
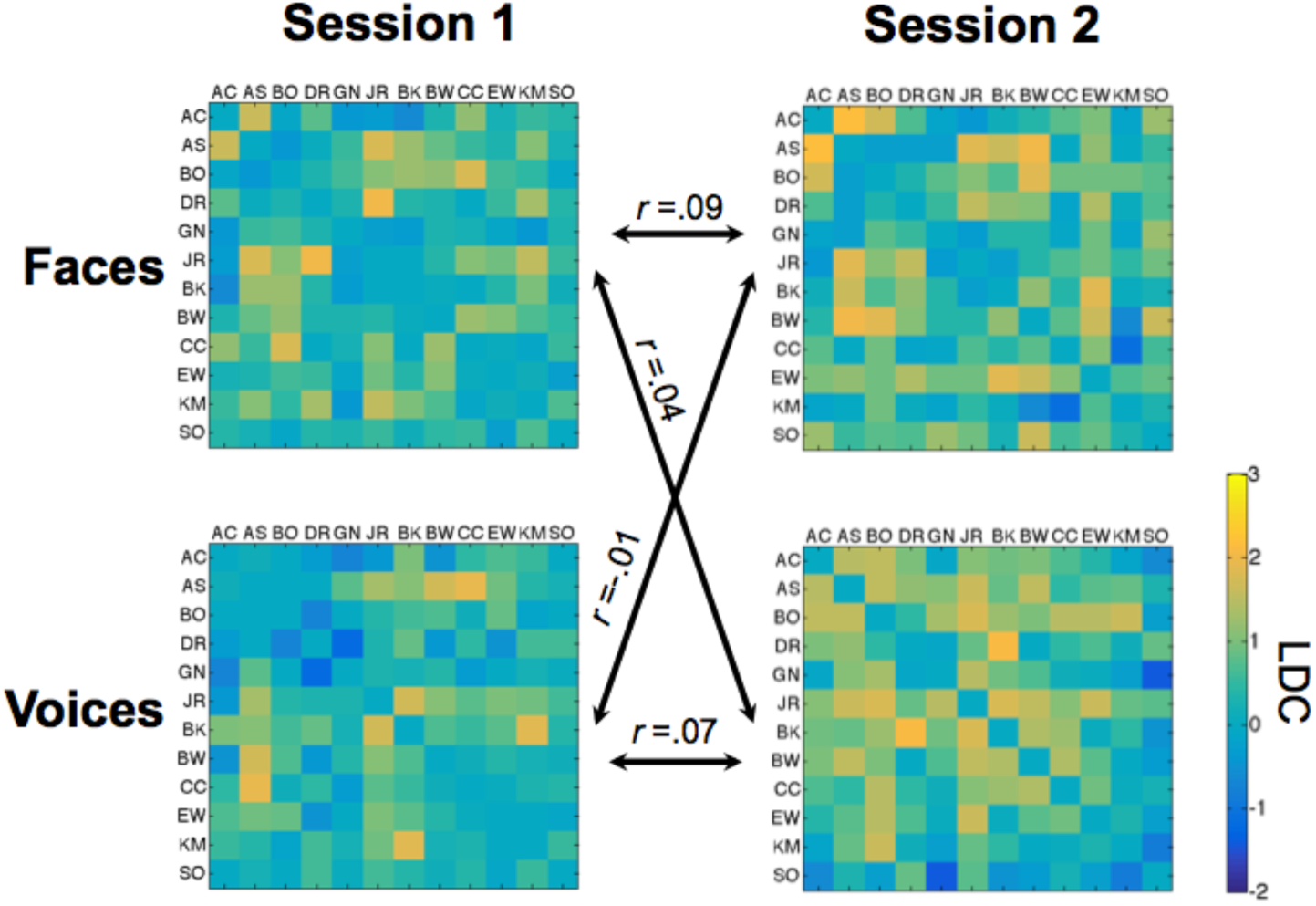
Representational distance matrix (RDM) comparisons across scanning sessions 1 and 2 in the rpSTS. Face and voice RDMs for the rpSTS were averaged across all 30 participants for illustration purposes. Each cell shows the discriminability of the brain activity patterns corresponding to a pair of identities (12 identities in total) computed using the linear discriminant contrast (LDC) and crossvalidating across data from three runs. Each matrix is symmetric around a diagonal of zeros. A value of zero or lower indicates no discriminability. For each participant we compared the representational geometry of the face and voice RDMs with the representational geometry in the RDM of the *other* modality (crossmodal comparisons) and in the RDM of the *same* modality (unimodal comparisons) using Pearson’s correlation. The figure shows Pearson’s correlations for all comparisons averaged across participants.

We predicted that face and voice RDMs would be correlated in ROIs that represent person-identity independently from modality. However, our results showed no significant correlations between face and voice RDMs in face-selective, voice-selective, or multimodal ROIs (Figure 3). It is possible that comparing RDM across different scanning sessions taking place on separate days did not allow us to detect subtle consistencies in the representational geometry for face-identities and voice-identities. To address this concern, we also compared face and voice RDMs within the same scanning session. However, we still found no significant correlations between face and voice RDMs. Therefore, using this method we found no evidence of modality-general person-identity representations in our ROIs.

We also predicted that there would be correlations between RDMs within the same modality in regions that represent only face-identity or only voice-identity. No correlations between face RDMs or between voice RDMs in any ROI were significant after correction for multiple comparisons.

### Analysis B: RSA investigating identity discriminability

Our second main analysis tested the generalisation of pattern discriminants from one modality to the other. More specifically, we computed crossmodal RDMs and we tested whether linear discriminants computed on pairs of faces could be used to discriminate between pairs of voices, and vice-versa. We also tested whether each ROI could discriminate between pairs of stimuli within the same modality. Mean LDC distances across all cells in crossmodal, face, and voice RDMs were compared against zero.

**Table 2.**
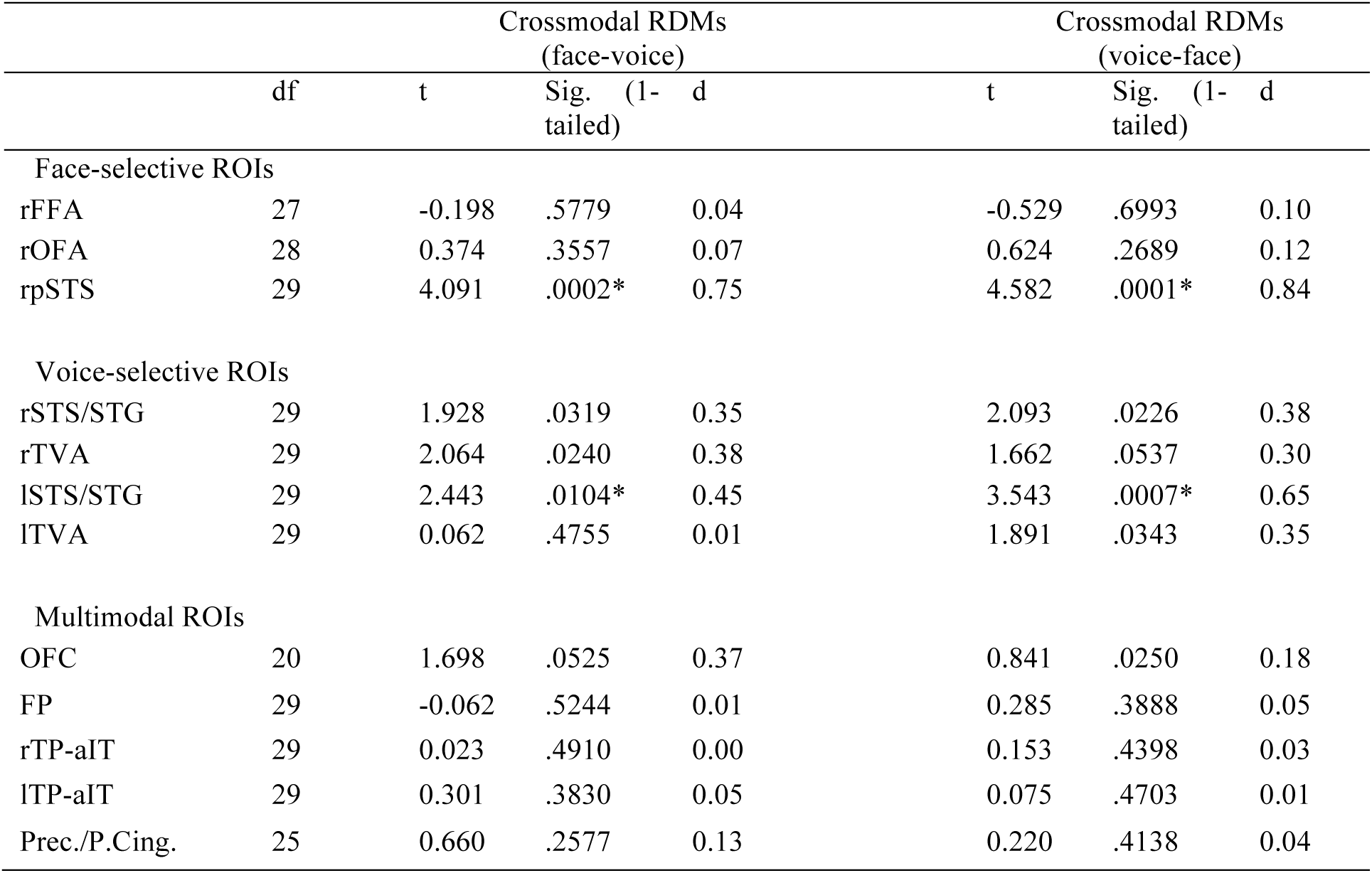
One-sample *t*-test results for mean LDC values in crossmodal RDMs. Stars represent statistical significance at *p*<.0150 (FDR corrected for all 48 comparisons in face, voice, and crossmodal RDMs). The rpSTS is classed as a face-selective ROI for consistency purposes only, but in fact it demonstrated multimodal properties.

**Table 3.**
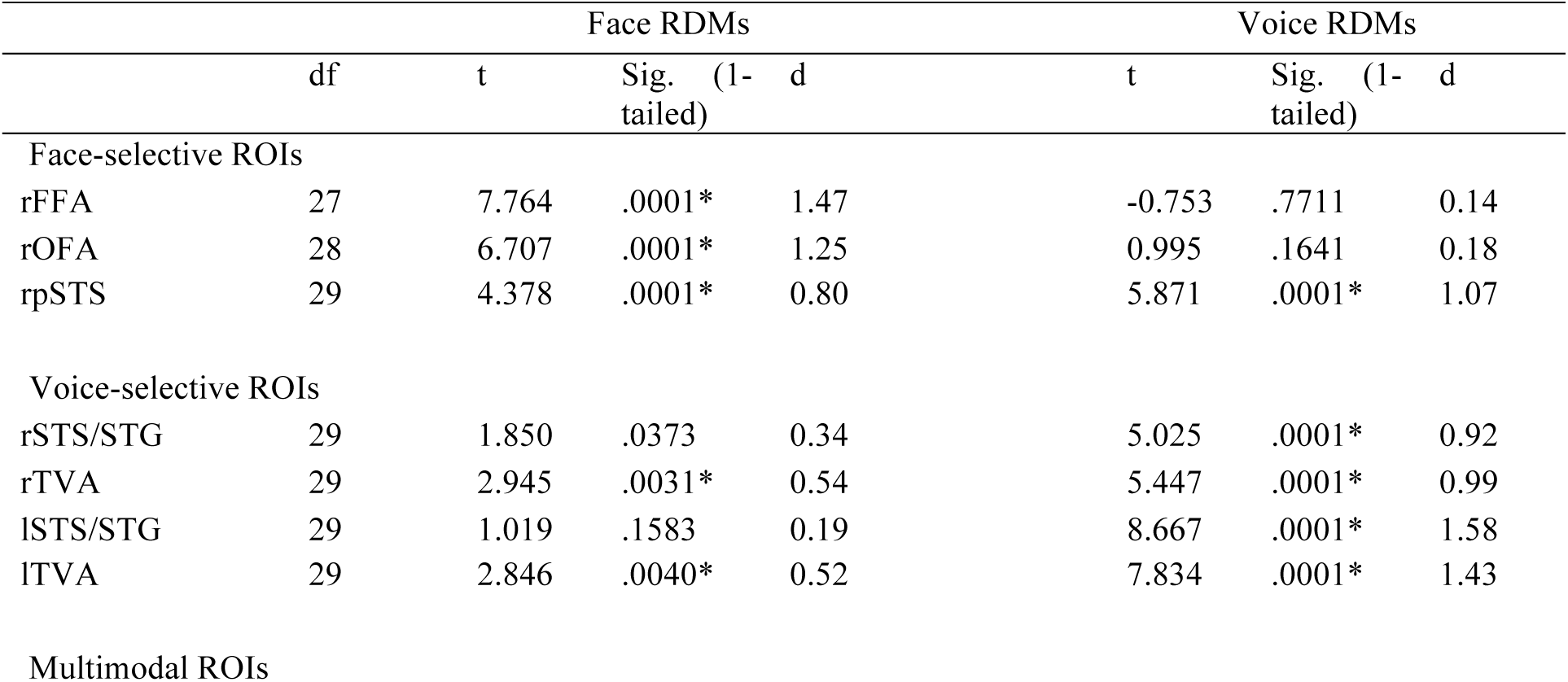
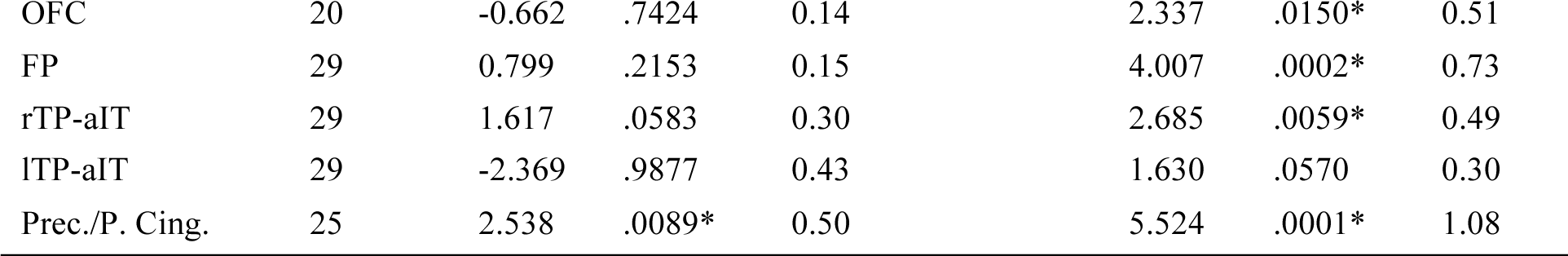
One-sample *t*-test results for mean LDC values in face and voice RDMs. Stars represent statistical significance at *p*≤.0150 (FDR corrected for all 48 comparisons in face, voice, and crossmodal RDMs). The rpSTS is classed as a face-selective ROI for consistency purposes only, but in fact it demonstrated multimodal properties.

We predicted that in brain regions with modality-general person identity representations the mean LDC values for crossmodal RDMs would be significantly greater than zero. Our results showed that mean LDC values in these RDMs were significantly greater than zero in the rpSTS, and in the voice-selective lSTS/STG (Figure 5; Table 2). These results show that the rpSTS could discriminate pairs of face-identities based on pattern discriminants computed from pairs of voice-identities (and vice-versa), and therefore appears to form modality-independent person-identity representations.

**Figure 5:**
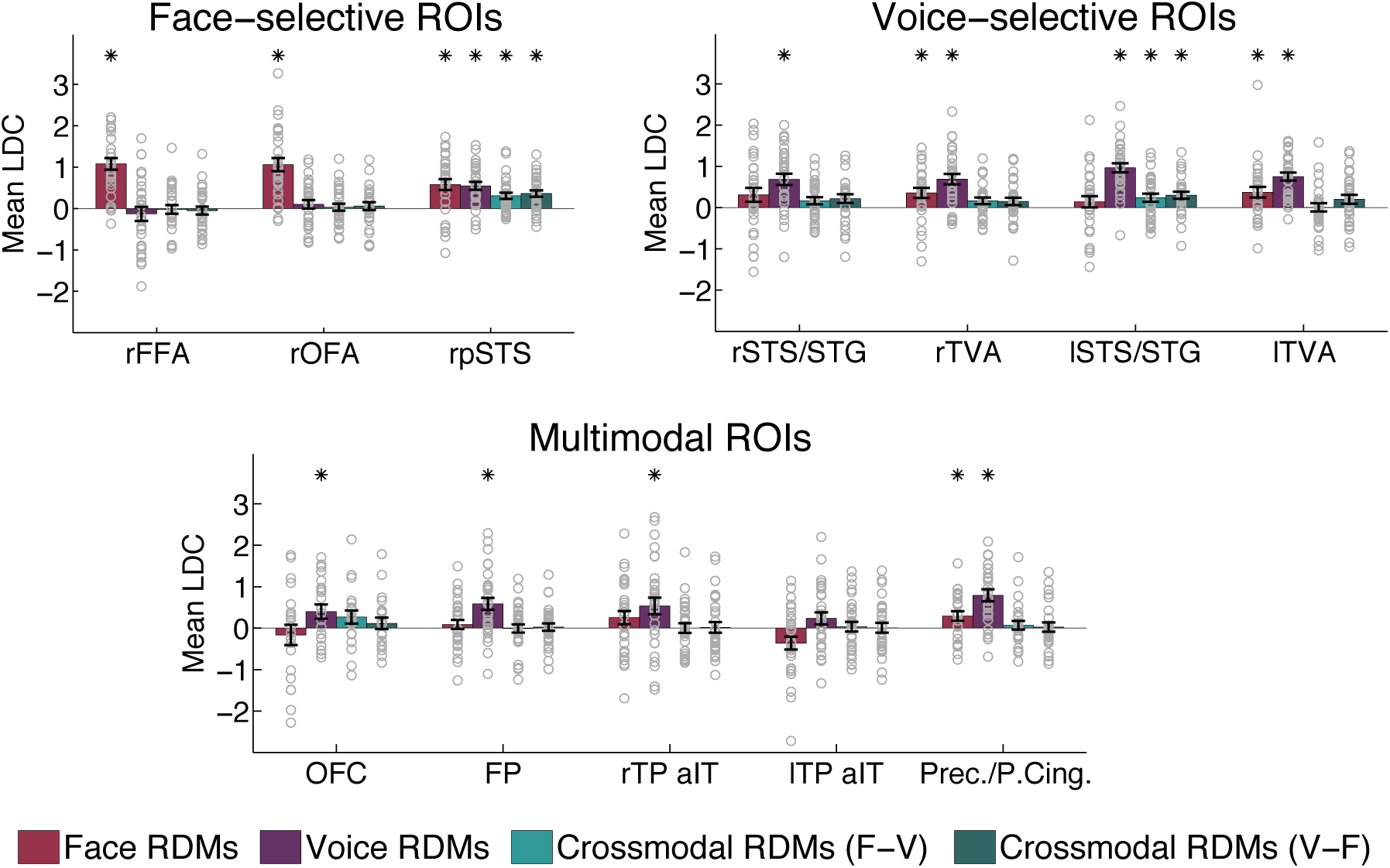
Results of RSA investigating identity discriminability. Mean LDC between identities in face RDMs, voice RDMs, and crossmodal RDMs in face-selective, voice-selective, and multimodal ROIs. There are two types of crossmodal RDMs: (a) face discriminant applied to voices (F-V), and (b) voice discriminant applied to faces (V-F). Bars show mean LDC values averaged across participants, error bars show standard error, and grey circles show mean LDC values for individual participants. We tested whether the mean LDC values were significantly greater than zero using one-sample t-tests across all 30 participants. Stars represent significant tests at *p*≤.0150 (FDR corrected for all 48 comparisons). These results show generalisation of the pattern discriminants from one modality to the other in the rpSTS and in the lSTS/STG. In addition, face-selective ROIs discriminate between face-identities, and voice-selective ROIs discriminate between voice-identities. Note that in this figure the rpSTS is classed as a face-selective ROI for consistency purposes only, but in fact it demonstrated multimodal properties.

We note that while the mean LDC values for crossmodal RDMs in the lSTS/STG were significant, the mean LDC value for face RDMs was not. While this result suggests that this region was able to discriminate identities based on crossmodal information, it is unlikely that a modality-general representation could exist without face-identity discrimination. Therefore, this result should be interpreted with caution. It is possible that in addition to the rpSTS, the lpSTS also contains a modality-general person-identity representation and it could be driving the positive result in the lSTS/STG. However, we were not able to test this because we could not localise the lpSTS in our participants using our face localiser.

We also predicted that mean LDC values for face RDMs and voice RDMs would be significantly greater than zero in ROIs that represent face-identity and voice-identity, respectively. We found that mean LDC values in face RDMs were significantly greater than zero in all ROIs originally defined as face-selective (rFFA, rOFA, rpSTS), in the TVAs, and in the multimodal Prec./P. Cing. (Figure 5; Table 3). These results show that all these regions could discriminate between face-identities. A follow up analysis in which all overlapping rpSTS voxels were removed from the rTVA showed that the significant result for faces in rTVA was driven by the rpSTS. Mean LDC values in voice RDMs were significantly greater than zero in all voice-selective ROIs (TVAs, STS/STG), in the rpSTS (originally defined as face-selective), and in the multimodal OFC, FP, rTP-aIT and Prec./P. Cing. (Figure 5; Table 3).

It is possible that the discrimination of identities in our ROIs was driven by different-gender identity pairs (female-male). To investigate this possibility, for each ROI and condition that showed mean LDC values significantly greater than zero (Figure 5 & Tables 2,3) and for each participant we compared the mean LDC values for different-gender identity pairs (calculated across 36 pairs) with the mean LDC values for same-gender identity pairs (calculated across 30 pairs: female-female & male-male) in each RDM (we used paired *t*-tests, and used FDR correction for all 19 comparisons). Results for the rpSTS showed no significant difference between the discriminability of different-gender and same-gender identity pairs for face, voice, or crossmodal RDMs (all *p*>.0533), demonstrating that person-identity discrimination in this region was not driven by discriminating gender. In contrast, mean LDC values for different-gender identity pairs were significantly higher than mean LDC values for same-gender identity pairs for face RDMs in the rFFA and rOFA (both *p* ≤.0010), and for voice RDMs in the bilateral TVAs and STS/STG (all *p*≤.0005), suggesting that gender contributed to the discrimination in these regions. However, mean LDC values for same-gender identity pairs were still significantly greater than zero (one-sample *t*-tests) for face RDMs in the rFFA and rOFA (both *p*<.0001) and for voice RDMs in the bilateral TVAs and STS/STG (all *p*≤.0239), suggesting that identity discrimination in these regions is not solely driven by differences in gender.

### Exploratory whole-brain searchlight analyses

Finally, we conducted additional exploratory searchlight analyses across the whole brain to determine whether there were brain regions with modality-general person-identity representations that are not included in our ROIs. The first searchlight analysis investigated correlations between face and voice RDMs across the whole brain, and we did not find any regions showing such correlations between face and voice representational geometries.

The second searchlight analysis investigated crossmodal generalization of discriminants for pairs of identities across the whole brain. We found a number of clusters in which the mean LDC in crossmodal RDMs was significantly greater than zero (FWE corrected threshold p ≤ .05), and below we report t-values and MNI coordinates for the peak grey matter voxels in each cluster. Anatomical labels for peak voxels are based on the Harvard-Oxford cortical and subcortical structural atlases. The results showed a large cluster (*k*=1927, *p*=.007) with peaks in the right putamen (*t*=4.33, x=21, y=20, z=−1), the left posterior middle temporal gyrus (*t*=4.04,x=−57, y=−19, z=−7), and the right precentral gyrus (*t*=3.89, x=54, y=8, z=32). Significant clusters were also found in the right paracingulate gyrus (k=1340, *p*=.003, *t*=4.34, x=6, y=47, z=23), in the left hippocampus (k=160, *p*=.017, *t*=4.45, x=−24, y=−37, z=2), in the right anterior supramarginal gyrus (k=84, *p*=.006, *t*=6.18, x=48, y=-22, z=38), in the left cuneal cortex (k=48, *p*=.036, *t*=3.99, x=−18, y=−76, z=29), and a cluster (*k*=100, *p*=.039) with peaks in the left temporooccipital middle temporal gyrus (*t*=3.58, x=−48, y=−46, z=5) and inferior lateral occipital cortex (*t*=3.45, x=−48, y=−67, z=8). Finally, we also found a significant cluster in the rpSTS at an uncorrected threshold of p ≤ .005 (k=592, *p*=.001, *t*=4.05, x=48, y=−49, z=11) that overlapped with our rpSTS ROI.

## Discussion

We show evidence of a modality-general person-identity representation in a multimodal region of the rpSTS, demonstrating that this region was able to discriminate familiar identities based on modality-general information in faces and voices. More specifically, the rpSTS could discriminate pattern estimates for pairs of face-identities based on linear discriminants computed from pattern estimates for pairs of voice-identities, and vice-versa. A crucial and novel aspect of our study is that we showed that the rpSTS not only discriminates between identities, but also generalises across multiple naturalistically varying face videos and voice recordings of the same identity. By always comparing pattern estimates across independent runs with different face and voice tokens for the same identities, we showed that the face- and voice-elicited person-identity representations in the rpSTS are stimuli-invariant. Invariant identity representations were also found for face-identities in face-selective regions (rFFA and rOFA) and for voice-identities in voice-selective regions (bilateral TVA and STS/STG). Finally, we did not find evidence of matching representational geometries for faces and voices, across or within modalities.

### A modality-general and invariant person-identity representation in the rpSTS

Our finding of a modality-general person-identity representation in a multimodal region of the rpSTS supports the Multimodal Processing Model, which proposes that face and voice information is integrated in multimodal brain regions (Ellis et al., 1997; Campanella and Belin, 2007). In contrast, we did not find support for the prediction from the Coupling of Face and Voice Processing Model (von Kriegstein et al., 2005; von Kriegstein and Giraud, 2006) that there would be a modality-general identity representation in face-selective regions of the fusiform gyrus.

The rpSTS has previously been associated with crossmodal representations of emotion from faces and voices and shows a preference for people-related stimuli regardless of modality (Watson et al., 2014a, 2014b). Furthermore, multiple studies have demonstrated overlap between face-selective and voice-selective regions in the rpSTS (Wright et al., 2003; Watson et al., 2014a; Deen et al., 2015; Anzellotti and Caramazza, 2017;). It has been proposed that the STS integrates person-specific patterns of movement from faces, voices, and bodies to assist in person-identity recognition (Yovel and O’Toole, 2016). It is possible that the intrinsic relationship between a person’s idiosyncratic facial movements and manner of speech contributes to the integration of face- and voice-identity information in the rpSTS.

Our finding of a modality-general identity representation in the rpSTS is also in agreement with two recent studies showing across-modality classification of pattern estimates for familiar faces and voices in the rpSTS (Anzellotti and Caramazza, 2017) and a more anterior part of the STS (Hasan et al., 2016). In contrast to these previous studies, we showed that the rpSTS also demonstrates face- and voice-elicited representations of person-identity that are invariant to different tokens of the same face and voice. The ability to “tell people together” by identifying different tokens of a face and voice as belonging to the same person is as important as the ability to “tell people apart” (i.e. discriminate between different people) (Burton, 2013; Anzellotti and Caramazza, 2014). Hasan et al. (2016) were unable to investigate invariant representations because they used a single face image and a single voice recording for each identity, which in turn were derived from the same original stimulus, making interpretation of their results difficult (Lavan, 2017). Anzellotti and Caramazza (2017) used two face and voice tokens for each identity but did not train and test their classifier on different tokens, and therefore did not demonstrate representations that were invariant to different tokens of the same face and voice in this study. Finally, in contrast to these previous studies, we used a larger set of identities and multiple naturalistically varying tokens in order to better capture the level of robust invariant recognition required in everyday life. While behavioural studies have shown the importance of within-person variability for recognition (Jenkins et al., 2011; Burton, 2013; Burton et al., 2016), this is rarely taken into account in neuroimaging experiments, which typically use highly similar or artificial stimuli for the same person.

### Invariant representations of face-identity and voice-identity

The face-selective rFFA and rOFA were able to discriminate between the faces of different people while also showing invariance to the different videos of each person’s face. This finding is in agreement with Anzellotti et al. (2014) and Guntupalli et al. (2017), who showed representations of face-identity in the FFA (and OFA, in Anzelotti et al., 2014) that generalise across different viewpoints of the face. However, in contrast with these studies, which used stimuli with low within-person variability, we show that representations in these regions generalise across highly variable face videos, and can thus discriminate between different face-*identities,* rather than between individual face *images.*

Voice-selective regions in STS/STG and the TVAs bilaterally could discriminate between different speakers while showing invariance to the different recordings of each voice. These findings are in line Formisano et al. (2008), who showed representations of speaker identity that generalise across utterances of different vowels in the lateral Heschl’s gyrus/sulcus and in the right STS. We extend this finding by showing that generalisation across different recordings of the same voice is possible even when using short sentences with variable speech content that were recorded in different settings.

We also found invariant discrimination of face- and voice-identity in a multimodal region in the precuneus/posterior cingulate. This region has been previously associated with the processing of familiar faces and voices (Shah et al., 2001), and has been found to discriminate between different face-identities (Visconti Di Oleggio Castello et al., 2017). Our results suggest that representations of faces and voices may be interspersed in this region, but are not shared across modalities. Finally, we showed invariant representations of voice-identity, but not face-identity, in the frontal pole, a region that has been previously associated with the processing of familiar voices (Nakamura et al., 2001). It should be noted that although we initially localised the frontal pole as a multimodal region, our results showed that it did not respond significantly to faces in the main experimental runs.

### Representational geometries

We did not find matching representational geometries across faces and voices in rpSTS despite finding crossmodal generalisation of the pattern discriminants. It is possible that all identities were equally distinct from each other within each modality (i.e. the nature of person-identity code in these regions does not result in variable representational distances between identities). In addition, the rpSTS shows both modality-specific and modality-general representations, and it is possible that the former had stronger influence on the representational geometry. Beauchamp et al. (2004) showed that the pSTS contains intermixed visual, auditory, and multisensory patches, and future studies could use higher-resolution neuroimaging methods to probe person-identity representations in this region.

In all other ROIs, we also did not find any evidence of stable representational geometries for face-identities or voice-identities only. Again, it could be that identities were equally distinct across from each other within each modality, or it could be that experimental conditions would need to be improved to obtain more reliable representational geometries.

### Anterior temporal lobe and searchlight results

We did not find evidence of face-, voice-, or person-identity representations in the anterior temporal lobe. This was surprising given that this region has been previously associated with the processing of person-identity (Ellis et al., 1989; Gainotti, 2011). The fact that our TP-aIT ROIs responded more to voices that to faces suggests that our multimodal region localiser was not optimal for detecting multimodal responses in the anterior temporal lobe. Moreover, our sequences were not tailored to detect fMRI responses in this region (Axelrod and Yovel, 2013), and therefore more research using specialised scanning parameters for the localisation of this region is warranted.

It is possible that modality-general representations exist outside face- and/or voice-selective regions, and our exploratory searchlight results revealed person-identity representations in the paracingulate gyrus, right insular cortex, left nucleus accumbens, left anterior postcentral gyrus, and left hippocampus. Quiroga et al. (2005, 2009) found that cells in the hippocampus (and also amygdala and entorhinal cortex) were highly responsive to specific identities, and responded to both the face and name of that person. It will be interesting to further probe the role of the hippocampus (and the other regions found during the searchlight analyses) in person-identity recognition.

## Conclusion

To conclude, we showed a modality-general person-identity representation that generalises across different, naturalistically varying face videos and voice recordings of the same person in a multimodal region of the rpSTS. This supports the Multimodal Processing Model for face and voice integration. We also found evidence of video-invariant face-identity representations in face-selective regions (rFFA, rOFA), and sound-invariant voice-identity representations in voice-selective regions (TVA, STS/STG). Future studies could focus on the nature and type of face and voice information that is represented in these different regions, and in how these representations are formed, both through development, and during the process of becoming familiar with someone.

## Acknowledgements

This work was supported by a research grant by the Leverhulme Trust (RPG-2014-392). We thank Matthew Longo for comments on a previous version of the manuscript, and Tiana Rakotonombana, Roxanne Zamyadi, and Rasanat Nawaz for help with preparing and piloting the stimuli. The authors declare no competing financial interests.

